# Reduced intra-tendinous sliding in Achilles tendinopathy during active plantarflexion regardless of horizontal foot position

**DOI:** 10.1101/2024.01.09.574669

**Authors:** Laura Lecompte, Marion Crouzier, Stijn Bogaerts, Lennart Scheys, Benedicte Vanwanseele

## Abstract

The Achilles tendon consists of three subtendons with the ability to slide relative to each other. As optimal intra-tendinous sliding is thought to reduce the overall stress in the tendon, alterations in sliding behavior could potentially play a role in the development of Achilles tendinopathy. The aims of this study were to investigate the difference in intra-tendinous sliding within the Achilles tendon during isometric contractions between asymptomatic controls and patients with Achilles tendinopathy and the effect of changing the horizontal foot position on intra-tendinous sliding in both groups. 29 participants (13 Achilles tendinopathy, 16 controls) performed isometric plantarflexion contractions at 60% of their maximal voluntary contraction (MVC), in toes-neutral, and at 30% MVC in toes-neutral, toes-in and toes-out positions during which ultrasound images were recorded. Intra-tendinous sliding was estimated as the superficial-to-middle and middle-to-deep relative displacement. Our results indicate that patients with Achilles tendinopathy present lower intra-tendinous sliding compared to asymptomatic controls. Regarding the horizontal foot position in both groups, the toes-out foot position resulted in increased sliding compared to both toes-neutral and toes-out foot position. We provided evidence that patients with Achilles tendinopathy show lower intra-tendinous sliding compared to asymptomatic controls. Since intra-tendinous sliding is a physiological feature of the Achilles tendon, the external foot position holds promise to increase sliding in patients with Achilles tendinopathy and promote healthy tendon behavior. Future research should investigate if implementing this external foot position in rehabilitation programs stimulates sliding within the Achilles tendon and improves clinical outcome.

## INTRODUCTION

The Achilles tendon (AT) is the largest, thickest and strongest tendon of the human body. The AT experiences substantial loads during everyday activities equivalent to 4 times body weight when walking and up to 12.5 times body weight when running and jumping^15^. Appropriate loading is crucial for a healthy tendon, however too much load leads to overuse marking the AT as one of the tendons most prone to injuries^29,34,35^. Achilles tendinopathy is a disabling musculoskeletal overuse disorder characterized by swelling and (activity-related) pain in the AT, with cumulative prevalence rates of 6% in physical exercise and reaching up 52% in specific athlete populations^32^. The prognosis of Achilles tendinopathy is often poor with high rates of recurrence ranging from 24 to 46%^36,52^. These findings highlight the limited understanding of the underlying mechanisms of Achilles tendinopathy. Therefore, gaining a deeper understanding of the mechanical behavior of both the healthy and tendinopathic AT is of main importance for improving both prevention and rehabilitation approaches.

Loading of the AT primarily comes from the 3 muscles of the triceps surae: Gastrocnemius medialis (GM), Gastrocnemius lateralis (GL) and Soleus (SOL). The subtendons of the two-headed superficial gastrocnemius fuse first around the middle region of the calf. More distally, the deeper SOL subtendon joins to form the free AT (i.e. free of muscle tissue). Because of this specific anatomical configuration, the AT is subjected to forces from 3 different muscles. How these forces are distributed among the heads of the triceps surae muscles influences the loading of the AT^24^. This muscle force distribution (also called muscle force sharing) has been regarded as a potential contributor to AT problems^6,28,40^. The subtendons do not only experience different loading, they can also slide relative to each other resulting in non-uniform motions (i.e. intra- tendinous sliding) during various tasks including passive, eccentric, isometric and dynamic (i.e. walking) exercises^1,5,46,47^. This ability of subtendons to slide relative to each other is thought to help distributing some of the tendon stretch, protecting the AT against excessive strain and reducing the overall stress^48^. There is consensus that aging leads to a more uniform AT deformation pattern, supported by findings in animals^48^, middle-aged individuals^47^ as well as elderly^8^. These changes may be due to age-related physiologic changes in the tendon such as increased collagen cross-linking^9^ or alterations in lubricin concentration^51^. In the context of pathology, only one pilot study has investigated the heterogeneous motions within the tendinopathic AT^10^. During sitting and standing heel raises, Couppé et al. (2020) indicated a tendency towards more uniform motions within the tendinopathic AT compared to the asymptomatic contralateral AT. However, compensatory mechanisms could be at play in the asymptomatic limb which might not provide an ideal reference for healthy tendon behavior comparison^26^. Moreover, the study by Couppé et al. focused on lower-intensity heel raises whereas Achilles tendinopathy often occur in active populations engaging in high-intensity activities (e.g. running). Whether this intra-tendinous sliding within the AT is caused by muscle force sharing strategies or rather by the tendon properties themselves is unclear. Only two studies examined muscle force sharing during isometric contractions and found alterations in patients with Achilles tendinopathy compared to asymptomatic controls^13,38^. However, the scarcity of available research makes it difficult to draw clear conclusions. Additional research is necessary to comprehensively understand the intra-tendinous sliding within the AT and its potential link to muscle force sharing in both healthy and tendinopathic individuals.

Interestingly, the intra-tendinous sliding that exists in the healthy AT is not an inalterable parameter, but can be modulated by changing the horizontal foot position, during isometric plantarflexion contractions^12^. It was found that the relative motions in the superficial part of the tendon tended to be larger during plantarflexion with the foot horizontally outwards rotated (toes-out position) compared to an internal horizontal rotation of the foot (toes-in position).

This suggests that altering the horizontal position of the foot may present an opportunity to enhance or diminish intra-tendinous sliding. However, it remains unclear whether the impact of foot position on intra-tendinous sliding is comparable between healthy individuals and patients with Achilles tendinopathy.

The purpose of the current study was to gain better insight in intra-tendinous sliding of the AT in Achilles tendinopathy by: (1) studying the difference in AT intra-tendinous between asymptomatic individuals and patients with Achilles tendinopathy during isometric plantarflexion at low and high intensities and (2) investigating whether the horizontal foot position affects AT intra-tendinous sliding in both groups. We hypothesized that intra- tendinous sliding would be lower in patients with Achilles tendinopathy when compared to asymptomatic controls, and that this would be explained by different force sharing strategies between the two groups. Furthermore, we expected the lower intra-tendinous sliding in Achilles tendinopathy to happen in all horizontal foot positions.

## MATERIAL AND METHODS

### Participants

For this study, 16 asymptomatic controls and 16 patients with Achilles tendinopathy (in both groups: 12 men and 4 women, Achilles tendinopathy: 9 unilateral, 7 bilateral) were recruited from a convenience sample. Prior to the start of the measurements, written informed consents were obtained from all participants and ethical approval for this study was provided by the Research Ethics Committee of KU Leuven (study number S65522). Inclusion criteria for the asymptomatic controls were being 18-60 years old and having no lower limb injury in the past 6 months. The same age-range was applied for patients with Achilles tendinopathy. The diagnosis of Achilles tendinopathy was drawn by a physiotherapist. Their inclusion was additionally based on the following criteria: i) experiencing pain for at least 2 months specifically located at the AT, ii) rated at least 2/10 on a pain scale at some time during most weeks, iii) the pain had to be exacerbated by activities such as running or jumping, iv) the pain had to be triggered by palpation, and v) their tendon must have shown ultrasound observations of hypo-echogenicity and/or thickening of the tendon. To obtain validated information regarding AT pain and disability, all patients completed the Victorian Institute of Sport Assessment—Achilles questionnaire (VISA-A)^43^. For all participants, physical activity level was estimated using the International Physical Activity Questionnaire (IPAQ), an evaluation tool of physical activity^11^. In the tendinopathic group, the affected leg (or in the case of bilateral Achilles tendinopathy: their most painful leg) was tested. Special attention was given to ensure that the groups were similar in terms of age, body mass index, and physical activity level. Overall, 3 patients with Achilles tendinopathy were excluded from analysis of which 2 because of low-quality data collection and 1 due to technical problems with the ultrasound analysis software resulting in a final sample size of 16 asymptomatic controls and 13 patients. Final participant characteristics are presented in Table 1.

**Table 1:** Analysed participant characteristics.

|  | **Tendinopathy (N = 13)** | **Controls (N = 16)** | ***p-value*** |
| --- | --- | --- | --- |
| **Age (years)** | 44 ± 13 | 39 ± 16 | 0.368 |
| **Height (m)** | 1.74 ± 0.10 | 1.72 ± 0.10 | 0.457 |
| **Weight (kg)** | 67 ± 14 | 67.5 ± 12 | 0.763 |
| **BMI (kg/m<sup>2</sup>)** | 23.9 ± 3.5 | 23.4 ± 4.8 | 0.852 |
| **Physical activity level** | 7582 ± 1538 | 5719 ± 3169 | 0.108 |
| **Sex** | 3 women , 10 men | 4 women, 12 men | - |
| **Measured leg** | 7 dominant , 6 non-dominant | 10 dominant , 6 non-dominant | - |

### Experimental design

Participants attended one laboratory session in the Movement & Posture Analysis Lab Leuven (Leuven, Belgium). First, while participants lie prone on a mattress, freehand 3D ultrasound was used to estimate the volume of GM, GL and SOL muscles. The freehand 3D ultrasound system consisted of a 15MHz 40-mm linear array ultrasound transducer (L15-7H40-A5, ArtUs EXT-1H system, UAB Telemed, Vilnius, Lithuania), equipped with a 3D printed 4-markers rigid body, tracked by a 3D motion capture system (V120:Trio tracking system, Optitrack, Corvallis, OR, USA). 2D B-mode ultrasound images and the 3D position of the transducer were synchronized by PlusServer software (public software library for ultrasound imaging research; v2.8.0; Kingston, Canada; Ungi et al., 2016) and recorded through 3D Slicer software using the SlicerIGT module (slicer.org; v4.10.1; Perth, Australia)^18, 53^.

Muscle volumes were estimated by scanning each muscle one by one, allowing for optimal position of both the leg and ultrasound probe for each muscle belly. For each muscle, depending on muscle size, 3 to 10 parallel scans were completed over the posterior lower leg. The ultrasound transducer was positioned transversally to the longitudinal axis of the tendon. To improve image quality and to avoid to put pressure on the skin that would have changed the muscle shape, a lot of gel was placed on the skin before every scan. Average total scanning time was about 50s for GL, 80s for GM and 120s for SOL.

Thereafter, participants were seated with the hip at 70° of flexion, the leg fully extended and the foot rigidly strapped in the isokinetic dynamometer (Biodex Medical Systems, Shirley, New York, USA). The ankle angle was fixed at 5° of plantarflexion. After familiarization with the plantarflexion task and a standardized warming-up, 4 isometric maximal voluntary contractions (MVC) of 5 seconds each, with 120 seconds rest in between, were executed. To ensure that each participant produced a true maximal contraction, the difference between the 2 highest contractions was calculated. If the difference was higher than 10%, an additional maximal contraction was performed. MVC’s were performed with the foot horizontally in the neutral position, i.e. the long axis of the foot aligned with the sagittal plane of the body.

Once the MVC’s were obtained, ramped (5s) submaximal isometric contractions at 30% and 60% of MVC were conducted in a randomized order. As heel raises, the most common exercise in the rehabilitation of Achilles tendinopathy, generally match an intensity of 22 to 30% of MVC, we wanted to test both this lower intensity and a higher intensity^16, 25, 42^. Contractions at 60% of MVC were conducted with the foot in the neutral position during which EMG-activity was measured. Contractions at 30% of MVC were conducted in toes-neutral, toes-in and toes- out positions, of which this foot rotation position was also applied in a randomized order. For the testing in toes-in, the foot was internally rotated by 30° from the long axis of the foot. For the testing in toes-out, the foot was externally rotated by 50° from the long axis of the foot. If participants were not capable of reaching this angle, their maximal range of motion was applied. In total, participants performed 16 contractions (4 repetitions for every individual task), while ultrasound images of the Achilles tendon were obtained. Fatigue was monitored by keeping a minimum resting time between contractions, and by further asking the participant if they felt like they could do the next contraction. Feedback of the torque was provided using visual feedback displayed on a monitor in front of the participant.

Myoelectrical activity was collected from the GM, GL and SOL. First, the participants tested leg was prepared by shaving and cleaning the skin to minimize the skin-electrode impedance and facilitate electrode fixation after which pairs of surface electrodes (Ag/AgCL electrodes, 10mm recording diameter, Ambu) were attached to the skin with a ∼ 20mm interelectrode distance (center-to-center). To minimize any potential cross talk during muscle activity and to ensure electrodes were positioned away from the borders of neighboring muscles and aligned with the direction of the fascicles, electrode location was first checked by B-mode Ultrasound. For GM and GL, the electrodes were placed on the middle line of the muscle belly at its two third distal. For SOL, the electrodes were placed below GM myotendinous junction. EMG signals were sampled at 2000 Hz via a MESPEC 8000 system (Mega Electronics Ltd., Finland).

During all isometric contractions, 2D-ultrasound images were acquired with the same 15 MHz 40-mm linear array ultrasound probe mentioned above. The transducer was placed longitudinally over the most distal part of the AT so that the calcaneal notch was visible to ensure repeatability of this positioning across participants. In addition, a piece of hypoechoic tape was attached to the skin at the level of the calcaneal notch to measure any possible displacement of the probe over the skin. The probe position was additionally drawn on the skin and for every contraction, the probe was placed on the marks and its orientation was adjusted consistently, i.e. so that the image showed superficial and deep tendon borders as clear as possible. When the rotation position of the foot was changed, care was taken to place the probe in the same position and orientation by using the skin marking. Ultrasound radio frequency data were collected at 70 frames/s using the Telemed Ultrasound Research Interface (ArtUs RF-Data Control, v1.4.4).

### Data processing

#### Speckle tracking analysis

A validated ultrasound speckle tracking algorithm (Matlab 2020b) was used to analyze the displacements of the superficial, middle and deep layers of the AT^14,45^. RF data were upsampled by a factor of 2 and 4 in the along-fiber (x) and transverse (y) directions^41^ to increase spatial density of correlation functions^30^. To track the displacement, a region of interest was selected on the ultrasound image, encompassing 3 rows by 13 columns of kernels, covering a 10 mm region along the tendon line of action (Figure 1). Frame-to-frame displacements were computed using a 2D normalized cross-correlation technique. Specifically, RF data within a kernel centered at the current node location were cross-correlated with RF data in a search window at the same location in the subsequent frame. A correlation threshold of r = 0.7 was used to discriminate valid displacement information^17,31^. These steps were repeated for all frames in a loading cycle to obtain the cumulative 2D displacement of each node. Motion tracking was also performed in the reverse direction by starting at the last frame and incrementing toward the first frame. The choice of data reduction and analysis parameters such as kernel size, window sizes and cross-correlation threshold was based on previous research, but additional options were included to optimize tracking quality^12^. In case the algorithm failed to identify tendon layer displacements, kernel and search window sizes were slightly changed until tracking worked. The range of kernel sizes used was between 1.2 and 2.2 mm (width) and 1.2 and 2 mm (height), while the search window ranged from 2.2 to 3.6 mm (width) and 1.2 to 3.6 mm (height). The algorithm calculated the displacement of every kernel and the average value of all kernels of the first, second and third row represented respectively the absolute displacement of the superficial, middle and deep layer. Intra-tendinous sliding was calculated as the differential displacement between the superficial and middle layer (i.e. superficial relative motions) and between the middle and deep layer (i.e. deep relative motions). Hence, negative values here represent respectively the superficial layer moving more than the middle layer and the middle layer moving more than the deep layer. The resulting tracking of each tendon layer was visually inspected by the investigator (LL). If the algorithm failed to track displacement, if image quality was poor or if the tracking didn’t seem to follow the actual displacement in an accurate way, these trials were excluded from analysis. On average, 3.3 out of the 4 trials per task could be included in the analysis (464 trials in total of which 10 were excluded because of poor quality and 72 because of failed tracking). The average value per task per subject was calculated and used in the analysis.

**Figure 1:**
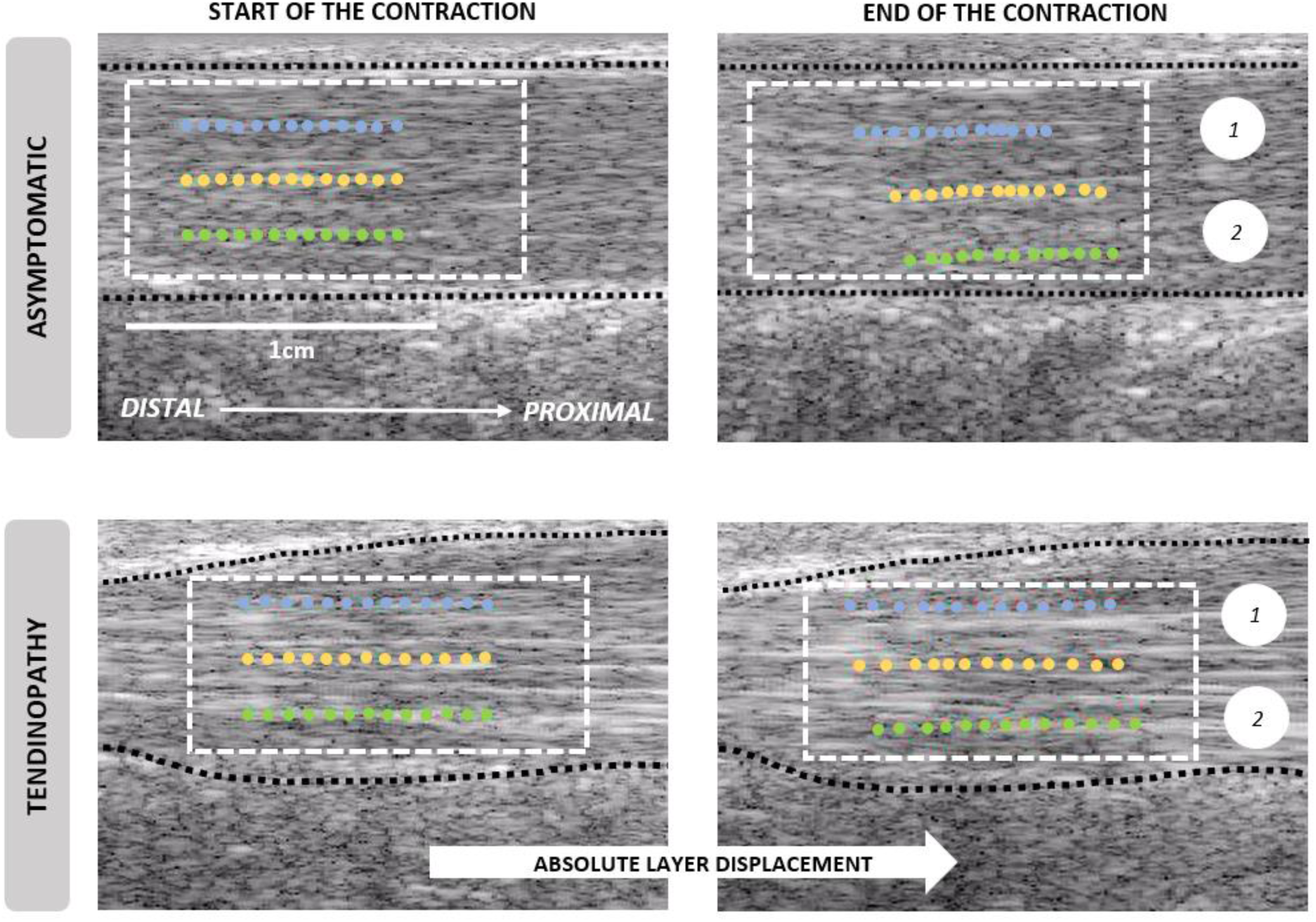
Speckle tracking algorithm of a tendinopathic and asymptomatic AT. The region of interest (white box) is selected wherein the kernels of the superficial (blue), middle (yellow) and green (deep) AT layers are tracked during ramped isometric contractions, allowing to track displacements. Dotted black lines represent the superficial and deep AT borders. 1 represents the superficial relative motions i.e. difference between the absolute displacement of superficial and middle layer), 2 represents the deep relative motions (i.e. difference between the absolute displacement of the middle and deep layer).

#### Reliability of intra-tendinous sliding

The within-session reliability of AT intra-tendinous sliding was evaluated. For each condition (i.e. 60% MVC neutral, 30% MVC neutral, 30% MVC toes-in and 30% MVC toes-out), all subjects with 3 or 4 good trials per condition were included. The intra-class correlation coefficient (ICC) and standard error of measurement (SEM = standard deviation * √ (1 – ICC)) were calculated. Our findings reveal an average ICC of 0.86 and SEM of 0.13mm, indicating a high level of reliability for our intra-tendinous sliding measures, within one measurement session. Further details on this reliability assessment can be found in supplementary material.

#### Torque and muscle activation

Both torque and EMG data were processed and analyzed using Matlab 2020b (MathWorks Inc., Natick, MA). Torque data was low-pass filtered at 10Hz. EMG signals were band-pass filtered (20–500 Hz) and full-wave rectified. Raw EMG signals were all visually inspected for electrical noise and movement artifacts. For the MVC trials, maximal torque and maximal EMG were calculated as the highest signal-value over a 300ms moving average window. The resulting highest EMG value over all trials was considered as the maximal RMS EMG value for further analysis. For the submaximal isometric force-matched tasks at 60% of MVC, the RMS EMG was calculated over 5s at the middle of the force plateau and normalized to that determined during the maximal isometric contractions.

#### Muscle volume

The volume reconstruction from the 2D ultrasound images was done in 3D Slicer using the Volume Reconstruction module^53^. Myotendinous junction positions were determined as the first slice (scanning from distal to proximal) in which the muscle was visible. Segmentations were then done by selecting the muscle area on the image in increments of 10 slices towards the proximal direction until the last slice where the muscle remained visible. In order to accurately capture the changing shape of the muscles, additional segmentations were performed near the origin and insertions of each muscle, as these areas exhibit a more variable muscle shape compared to the tubular muscle belly. Using the Fill In Between Slice function in 3D slicer, segmentations were combined and muscle volume was reconstructed. Segmented slices of each muscle were checked visually by the investigator and if necessary, additional segmentations were performed to refine and improve the muscle volume reconstruction.

#### Index of force

Based on previous research, we considered that the difference in force produced by synergist muscles during isometric plantarflexion tasks depends mainly on their difference in activation and volume^13, 27^. An index of force (in arbitrary unit) was, therefore, calculated as the product of muscle activation (normalized RMS EMG) and volume (cm^3^) for the contractions at 60% of MVC with toes-neutral. Ratio’s were further calculated to represent the distribution of force among the triceps surae (TS), and calculated as each individual muscle force index over the sum of GM, GL and SOL force indices. They are reported as GM/TS, GL/TS and SOL/TS respectively.

### Statistics

Distribution of the data was checked for normality using a Shapiro–Wilk test (IBM SPSS Statistics 28.0). Demographic characteristics of participants, tendon thickness and MVC were compared between groups using independent t-tests. To determine whether muscle force-sharing was different between groups, a two-way repeated measures ANOVA was conducted (within-subject factor: muscle force index [GM/TS, GL/TS and SOL/TS]; between-subject factor: group [asymptomatic control and Achilles tendinopathy]). To determine whether the intensity affected the intra-tendinous sliding differently between groups and across tendon regions, a three-way repeated measures ANOVA was conducted (within subject-factors: AT region [superficial and deep] and intensity [30% and 60% of MVC]; between-subject factor: group). Similarly, another three-way repeated measures ANOVA was conducted to determine whether the horizontal foot position affected the intra-tendinous sliding differently between groups across tendon regions (within subject-factors: AT region and foot position [toes-neutral, toes-in and toes-out]; between-subject factor: group). When appropriate, post hoc analysis were performed using the Bonferroni test. The level of significance was set at p ≤ 0.05. The eta- squared (ղ2) values were reported as measures of effect size, with 0.01, 0.06, and 0.14 as small, medium, and large effects, respectively. Data are reported as mean ± standard deviation.

## RESULTS

### Muscle and tendon characteristics

The AT in the tendinopathic group was significantly thicker compared to the asymptomatic control group (tendinopathy: 8.6 ± 1.8 mm, controls: 6.1 ± 0.8 mm, p < 0.001). No significant between-groups differences were found on the muscle-force sharing (all p > 0.160). Data are reported in Table 2.

**Table 2:** Muscle and force related characteristics in the tendinopathy and control group.

|  | Control | Tendinopathy |
| --- | --- | --- |
| *Torque during maximal voluntary contraction in plantarflexion (Nm)* |  |  |
| | $167.2 \pm 54.7$ | $142.1 \pm 54.7$ |
| *Muscle volume (cm<sup>3</sup>)* |  |  |
| GM | $245 \pm 60.5$ | $241 \pm 54.5$ |
| GL | $149 \pm 34.7$ | $145 \pm 35.9$ |
| SOL | $482 \pm 77.1$ | $491 \pm 99.7$ |
| *Muscle activation during contractions at 60% MVC (% of RMS EMG<sub>max</sub>)* |  |  |
| GM | $41.9 \pm 5.9$ | $38.6 \pm 11.8$ |
| GL | $34.7 \pm 9.1$ | $36.8 \pm 13.9$ |
| SOL | $45.1 \pm 15.0$ | $42.9 \pm 17.8$ |
| *Force index at 60% MVC (arbitrary unit)* |  |  |
| GM | $10.1 \pm 2.9$ | $9.7 \pm 3.8$ |
| GL | $5.2 \pm 2.1$ | $5.6 \pm 2.6$ |
| SOL | $21.9 \pm 7.2$ | $20.9 \pm 9.1$ |
| *Force index ratio at 60% MVC (%)* |  |  |
| GM/TS | $27.5 \pm 4.3$ | $27.7 \pm 10.2$ |
| GL/TS | $14.0 \pm 4.8$ | $15.6 \pm 5.0$ |
| SOL/TS | $58.5 \pm 7.5$ | $56.7 \pm 11.3$ |

### Achilles tendon displacement

For a matter of clarity, only relative motions between AT layers (deep-to-middle and middle-to-superficial) are reported. Absolute values of all AT layers displacement can be found in supplementary material.

#### Effect of intensity of contraction on intra-tendinous sliding

Main effects of *group* [F(1,27) = 10.333, p = 0.002, ղ^2^ = 0.28], *layer* [F(1,27) = 19.571, p < 0.001, ղ^2^ = 0.42] and *intensity* [F(1,27) = 8.717, p = 0.006, ղ^2^ = 0.24] were found on the relative motions within the AT. This demonstrated that i) people with tendinopathy generally present lower intra-tendinous sliding compared to asymptomatic controls (tendinopathy: 0.50 ± 0.09 mm, 95% CI [0.31 to 0.68]; control: 0.89 ± 0.08 mm, 95% CI [0.72 to 1.05]), that ii) the deep relative motions are larger than the superficial relative motions (deep: 0.95 ± 0.10 mm, 95% CI [0.75 to 1.15]; superficial 0.43 ± 0.07 mm, 95% CI [0.29 to 0.58]) and that iii) intra-tendinous sliding is larger at higher than at lower intensities of contraction (high: 0.77 ± 0.06 mm, 95% CI [0.64 to 0.90]; low: 0.61 ± 0.07 mm, 95% CI [0.47 to 0.75]). Interestingly, the layer * group interaction effect (p = 0.060, ղ^2^ = 0.13) showed a trend towards the fact that superficial relative motions were lower in the tendinopathy group compared to asymptomatic controls, regardless of intensity (see figure 2). The effect of intensity on intra-tendinous sliding in both groups can be seen on Figure 2.

**Figure 2:**
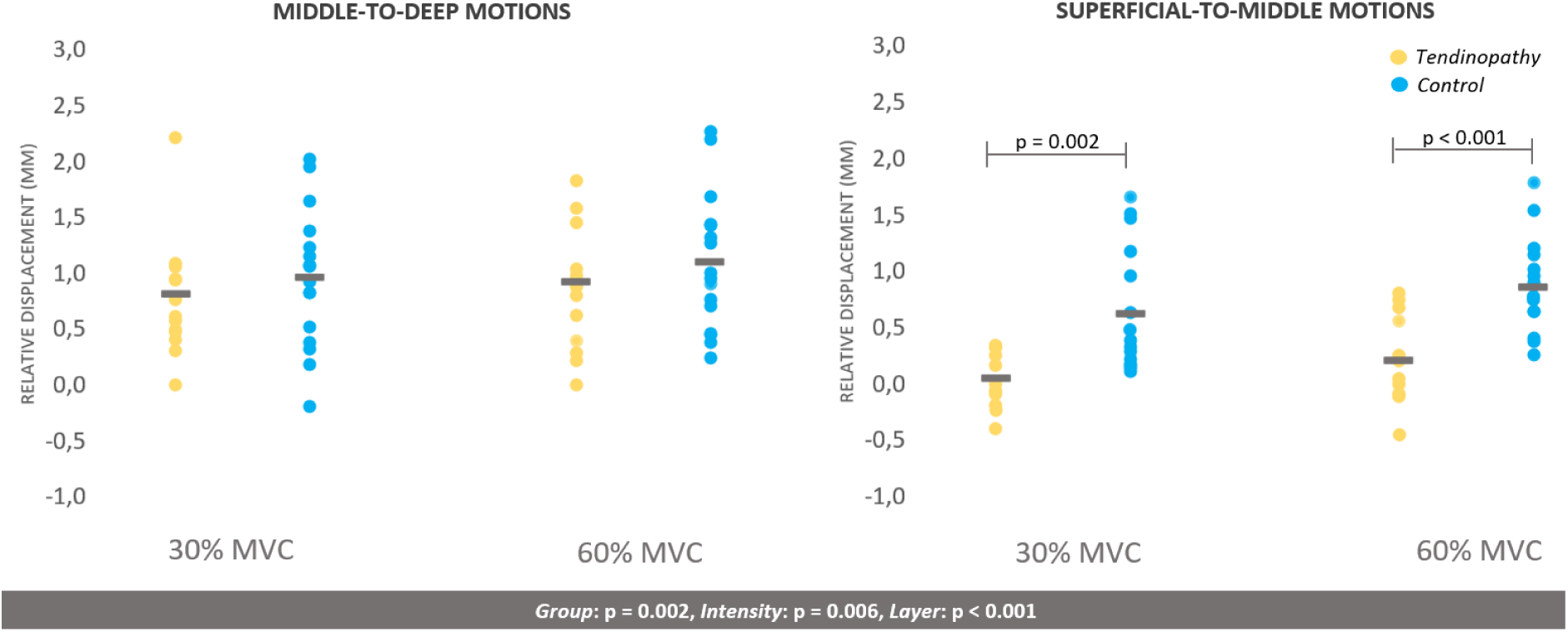
Effect of intensity on intra-tendinous sliding: deep (i.e. middle-to-deep) and superficial (superficial-to-middle) relative motions at 60% and 30% of MVC in the tendinopathy group (yellow) and asymptomatic control group (blue).

#### Effect of horizontal foot position on intra-tendinous sliding

Main effects of *group* [F(1,27) = 5.879, p = 0.022, ղ^2^ = 0.18], *layer* [F(1,27) = 34.405, p < 0.001, ղ^2^ = 0.56] and *foot position* [F(2,54) = 12.338, p < 0.001, ղ^2^ = 0.31] were found on the relative motions within the AT. This demonstrated that i) people with tendinopathy generally present lower intra-tendinous sliding compared to asymptomatic controls (tendinopathy: 0.50 ± 0.09 mm, 95% CI [0.31 to 0.68]; control: 0.80 ± 0.08 mm, 95% CI [0.63 to 0.96]), that ii) the deep relative motions are larger than the superficial relative motions (deep: 0.97 ± 0.10 mm, 95% CI [0.77 to 1.18]; superficial 0.33 ± 0.06 mm, 95% CI [0.21 to 0.44]) and that iii) that intra-tendinous sliding is larger in more toes-out than toes-in position (toes-neutral: 0.61 ± 0.07, 95% CI [0.47 to 0.75]; toes-in: 0.57 ± 0.07, 95% CI [0.43 to 0.70]; toes-out: 0.77 ± 0.06, 95% CI [0.64 to 0.90]. No group interaction effects were found. The effect of horizontal foot position on intra-tendinous sliding in both groups can be seen on Figure 3.

**Figure 3:**
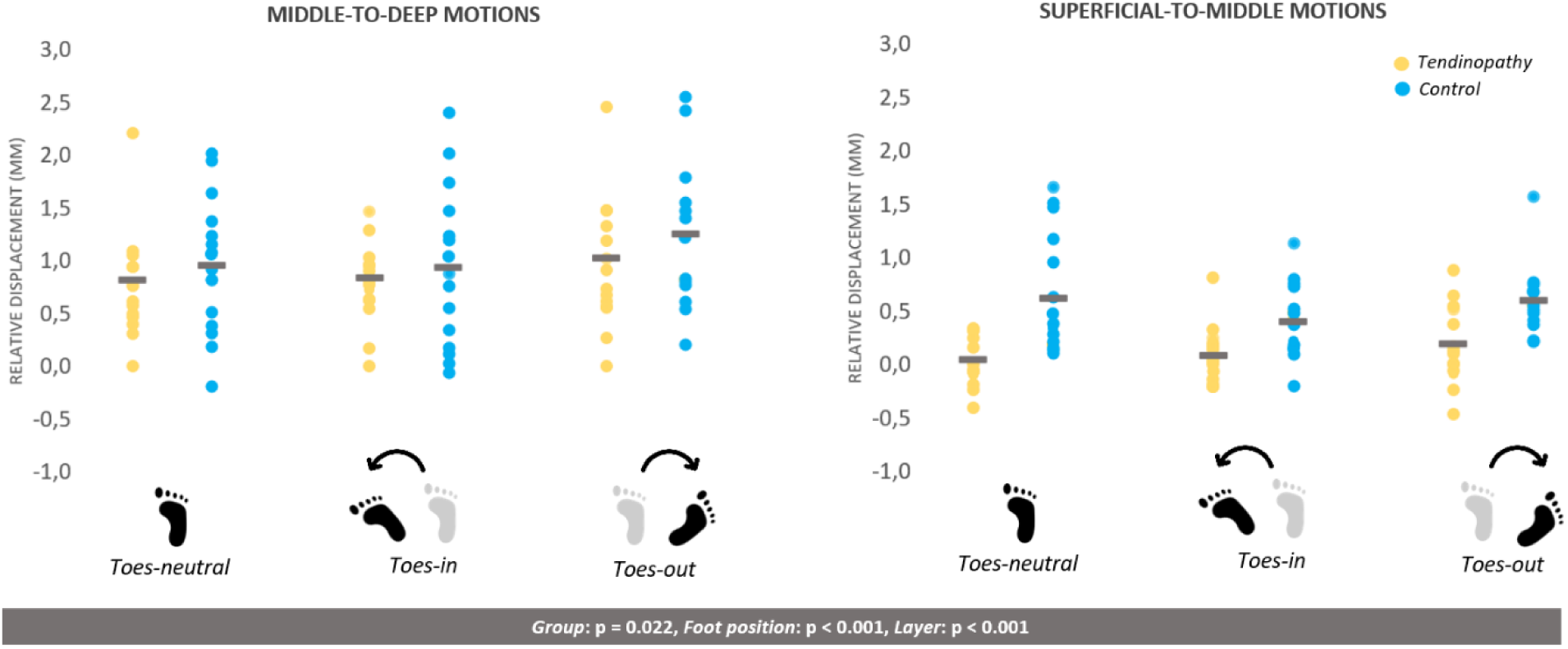
Effect of horizontal foot position on intra-tendinous sliding: deep (i.e. middle-to-deep) and superficial (superficial-to-middle) relative motions at 30% of MVC with toes-neutral, toes-in and toes- out in the tendinopathy group (yellow) and asymptomatic control group (blue).

## DISCUSSION

The primary aim of this study was to document the possible difference in intra-tendinous sliding within the AT between asymptomatic individuals and patients with Achilles tendinopathy. As hypothesized, we found a notable reduction in intra-tendinous sliding within the tendinopathic AT compared to asymptomatic controls. In other words, displacements were more homogeneous in the presence of tendinopathy compared to an asymptomatic tendon, despite having the same muscle force sharing. Further, we observed that both intensity and horizontal foot position played a significant role in influencing the sliding pattern, resulting in enhanced intra-tendinous sliding when increasing the intensity as well as with the foot turning outwards.

To the best of our knowledge, our findings regarding reduced intra-tendinous sliding in Achilles tendinopathy can only be compared to the results obtained by Couppé et al. (2020)^10^. In line with our research, they observed diminished superficial-to-deep displacement in the affected leg of individuals with unilateral tendinopathy, as compared to the asymptomatic leg, during both standing and sitting heel raises. Couppé et al. reported sliding values of 0.36 ± 0.20 mm during standing heel raises in the tendinopathic leg. Our superficial-to-deep displacements were higher, amounting to 0.87 ± 0.57 and 1.13 ± 0.57 mm during contractions at 30% and 60% of MVC respectively, which is likely due to the higher intensity of the tasks assessed in our study. This aligns with the normalized triceps surae EMG RMS value of 23.3% during standing heel raises reported by Hébert-Losier et al. (2012) compared to 40.6% found in our study during isometric plantarflexion at 60% of MVC^25^. This difference in contraction intensity may explain the higher displacements observed at the tendon level.

Differences in intra-tendinous sliding between the groups were predominantly seen in the superficial-to-mid region with almost no or even a negative relative displacement seen in the tendinopathy group. Interestingly, in 11 of the 13 patients, a dark hypo-echoic area primarily located in the superficial region was observed on the AT ultrasound images that were acquired to evaluate the inclusion criteria (at the midportion level). This darker area indicates a pathological change in tissue density, mainly characterized by a loss of collagen fibril integrity, fluid accumulation and the development of scar tissue^3^. Therefore, these differences in intra- tendinous sliding (captured more distally, e.g. the probe was just above the calcaneal notch) between the groups in the superficial-to-mid region may be linked to the presence of the tendinopathic area more superficially. In half of the patients, a higher displacement in the superficial layer compared to the middle layer was observed, a pattern which was not observed in any of the asymptomatic controls. Bogaerts et al. (2018) also found this “reversed” pattern, i.e. higher displacement in the superficial layer compared to deeper in the tendon, in combination with a distinct hypo-echoic area in the superficial layer in 2 of the 3 most pathological tendons in a pilot study^4^. These findings are also consistent with some evidence suggesting that collagen proliferation and scar tissue formation could explain lower sliding in Achilles tendinopathy^2, 37^. Since Achilles tendinopathy is typically an overuse disorder, repetitive overloading without adequate healing could lead to the accumulation of micro- ruptures, resulting in the formation of scar tissue. This process could contribute to the limited inter-fascicular sliding^50^. Another potential factor is the generation of cross-links due to the accumulation of advanced glycation end-products that occurs with aging^39^. These cross-links lead to morphological changes in the collagen network, ultimately affecting the biomechanical properties of the tendon tissue^23, 44^. In animal models, it is already proven that glycation reduces inter-fibrillar sliding capacity^19, 22, 33^. Taken together, this evidence supports the fact that the presence of a hypo-echoic area and the formation of interfascicular scar tissue could be closely linked to the altered intra-tendinous sliding within the tendinopathic AT. However, the exact cause-and-effect relationship remains unknown.

In our study, we did not find any differences in the triceps surae force indices between tendinopathy patients and asymptomatic controls, which is not consistent with previous findings reporting different contribution of GL to the triceps surae force^13^ or different SOL activation^40^. It is important to note that there were some differences in the studies design (e.g. position of the participants, intensities of contraction, physical activity level, symptom duration, etc.). These varying results highlight the limited knowledge we currently have regarding muscle force-sharing in patients with Achilles tendinopathy. Our study was the first to couple muscle force data with the free Achilles tendon internal behavior. Despite not finding differences in muscle forces between groups, we did observe important differences in their intra-tendinous sliding behaviors. This provides support to the fact that, changes occurring at the tendon level (for example, interfascicular scar tissue) rather than changes in muscle force-sharing have an important role in these altered AT motions.

Similar to healthy individuals, the horizontal foot position during plantarflexion has an impact on intra-tendinous sliding for patients with Achilles tendinopathy, especially as sliding behavior can be increased when a plantarflexion is performed with the toes-out. This opens new perspectives in the field of rehabilitation. Based on our findings, one possible way to promote healthy tendon behavior could be by adjusting the horizontal foot position, turned outwards, during rehabilitation exercises. Findings of Thorpe et al. suggest a role for the interfascicular matrix in facilitating sliding between fascicles allowing large extensions within the energy-storing tendons, without exposing them to excessive strains^49^. Further studies should explore the effects of implementing exercises in different horizontal foot positions during rehabilitation and refine rehabilitation approaches for Achilles tendinopathy.

This study is not without certain limitations. First, the complex 3D structure of the AT cannot be fully captured by 2D ultrasound imaging, which may result in some out-of-plane motions. Secondly, small changes in probe position might have had an impact on displacement tracking and impacted the comparison between conditions (i.e. intensities and foot positions). To minimize variability, we recorded 4 ultrasound videos per condition, utilized tape and markings on the skin for consistent probe placement and visually checked the ultrasound image for similarity. The within-session reproducibility further supports the consistency between trial acquisitions. Next, 3 patients with Achilles tendinopathy had to be excluded from analysis due to technical problems resulting in a smaller tendinopathy group compared to the control group. Since the two groups still showed similar demographic characteristics (age, body mass index and physical activity level), this should not have impacted the reported results. It is also important to note that there are no non-invasive techniques to directly measure individual muscle force. Thus, our study used an indirect method to estimate the force by multiplying muscle activation and muscle volume. Lastly, the cross-sectional design limits the ability to establish causality between intra-tendinous sliding alteration and Achilles tendinopathy. Knowing if the loss of intra-tendinous sliding is a cause or a consequence of the pathology would allow for better focus on either prevention strategies or treatment approaches.

## CONCLUSION

We provided evidence that patients with Achilles tendinopathy show reduced intra-tendinous sliding compared to asymptomatic controls. As the toes-out foot position generates higher intra- tendinous sliding in patients with Achilles tendinopathy compared to neutral position, future research should investigate if this external foot position can help to restore physiological tendon sliding and if its implementation in Achilles tendinopathy rehabilitation programs would fasten recovery.

