## Supplementary Material for "Reduced intra-tendinous sliding in Achilles tendinopathy during active plantarflexion regardless of horizontal foot position"

Absolute displacement of the Achilles tendon layers in the Achilles tendinopathy group

|  | 60% OF MVC | | | 30% OF MVC | | | | | | | | |
| --- | --- | --- | --- | --- | --- | --- | --- | --- | --- | --- | --- | --- |
|  | NEUTRAL | | | NEUTRAL | | | IN | | | OUT | | |
| *Subject number* | bottom | mid | up | bottom | mid | up | bottom | mid | up | bottom | mid | up |
| *3* | 8,69 | 7,65 | 6,90 | 3,04 | 2,55 | 2,21 | 2,43 | 1,52 | 1,32 | 2,11 | 1,54 | 1,44 |
| *4* | 2,95 | 1,50 | 0,70 | 2,43 | 0,22 | 0,62 | 0,88 | 0,25 | 0,11 | 3,25 | 0,79 | 0,85 |
| *5* | 9,56 | 7,73 | 7,48 | 6,21 | 5,16 | 4,91 | 7,50 | 6,47 | 5,65 | 9,14 | 7,66 | 6,78 |
| *6* | 1,17 | 0,95 | 0,71 | 1,20 | 0,80 | 0,87 | 0,71 | 0,54 | 0,52 | 1,88 | 1,61 | 1,23 |
| *7* | 4,29 | 3,31 | 3,42 | 3,36 | 2,59 | 2,65 | 3,10 | 2,47 | 2,60 | 3,39 | 2,20 | 2,43 |
| *10* | 2,61 | 1,70 | 1,80 | 1,31 | 0,74 | 0,93 | 1,79 | 1,02 | 1,20 | 2,19 | 1,46 | 1,29 |
| *13* | 5,29 | 4,41 | 4,37 | 3,66 | 2,71 | 2,77 | 2,89 | 1,93 | 1,88 | 3,13 | 1,79 | 1,67 |
| *17* | 2,27 | 1,64 | 1,75 | 2,19 | 1,72 | 1,40 | 1,37 | 0,55 | 0,76 | 4,21 | 3,30 | 2,75 |
| *18* | 1,52 | 0,72 | 0,81 | 1,48 | 0,86 | 0,95 | 1,52 | 0,79 | 0,84 | 1,02 | 0,46 | 0,54 |
| *19* | 3,36 | 2,42 | 2,21 | 1,20 | 0,90 | 0,73 | 0,97 | 0,42 | 0,25 | 1,68 | 1,00 | 0,99 |
| *20* | 3,18 | 1,60 | 2,04 | 2,52 | 1,43 | 1,66 | 2,47 | 1,18 | 1,39 | 2,44 | 0,95 | 1,41 |
| *22* | 2,77 | 2,48 | 1,81 | 2,35 | 1,41 | 1,08 | 2,10 | 1,15 | 0,82 | 2,01 | 1,39 | 0,74 |
| *24* | 2,15 | 1,76 | 1,20 | 1,88 | 1,11 | 0,78 | 3,50 | 2,03 | 1,78 | 2,61 | 1,58 | 1,06 |
| Average | 3,83 | 2,91 | 2,71 | 2,53 | 1,71 | 1,66 | 2,40 | 1,56 | 1,47 | 3,00 | 1,98 | 1,78 |
| St. Dev. | 2,59 | 2,33 | 2,25 | 1,37 | 1,30 | 1,22 | 1,77 | 1,62 | 1,44 | 2,02 | 1,85 | 1,63 |

Absolute displacement of the Achilles tendon layers in the asymptomatic control group

|  | 60% OF MVC | | | 30% OF MVC | | | | | | | | |
| --- | --- | --- | --- | --- | --- | --- | --- | --- | --- | --- | --- | --- |
|  | NEUTRAL | | | NEUTRAL | | | IN | | | OUT | | |
| *Subject number* | bottom | mid | up | bottom | mid | up | bottom | mid | up | bottom | mid | up |
| 8 | 3,19 | 1,76 | 0,85 | 3,31 | 2,48 | 2,37 | 3,21 | 2,66 | 2,24 | 2,98 | 2,16 | 1,40 |
| 9 | 11,42 | 9,14 | 7,60 | 11,85 | 9,89 | 8,93 | 11,89 | 9,48 | 8,74 | 10,23 | 7,67 | 6,90 |
| 12 | 3,83 | 2,87 | 2,60 | 4,70 | 3,46 | 3,28 | 2,95 | 1,75 | 1,37 | 4,20 | 2,79 | 2,38 |
| 14 | 4,20 | 2,76 | 1,80 | 3,08 | 1,06 | 0,77 | 2,51 | 1,74 | 1,25 | 3,94 | 2,46 | 1,77 |
| 15 | 3,61 | 3,22 | 2,02 | 1,51 | 1,12 | 0,90 | 1,37 | 1,25 | 0,72 | 2,45 | 1,90 | 1,42 |
| 16 | 1,94 | 1,70 | 1,04 | 0,86 | 0,53 | 0,39 | 0,58 | 0,55 | 0,36 | 2,54 | 1,93 | 1,37 |
| 21 | 5,91 | 4,22 | 3,84 | 4,42 | 3,04 | 2,64 | 4,36 | 2,88 | 2,39 | 3,76 | 1,96 | 1,44 |
| 23 | 4,07 | 2,74 | 2,09 | 4,74 | 3,09 | 1,91 | 3,30 | 1,55 | 1,75 | 5,50 | 3,06 | 2,83 |
| 25 | 3,68 | 3,22 | 2,19 | 0,97 | 1,16 | 0,83 | 1,25 | 1,31 | 1,09 | 1,70 | 1,48 | 0,90 |
| 26 | 2,25 | 1,80 | 1,39 | 0,86 | 0,67 | 0,51 | 1,01 | 0,83 | 0,73 | 3,16 | 2,32 | 2,10 |
| 27 | 4,83 | 4,12 | 3,24 | 3,98 | 3,46 | 1,94 | 2,92 | 2,57 | 1,77 | 3,85 | 3,07 | 2,70 |
| 28 | 4,83 | 4,05 | 2,90 | 4,84 | 3,91 | 2,43 | 5,84 | 3,82 | 3,05 | 4,97 | 3,41 | 2,85 |
| 29 | 7,50 | 6,59 | 4,80 | 5,44 | 4,61 | 2,94 | 5,32 | 4,43 | 3,29 | 5,50 | 4,70 | 3,12 |
| 30 | 4,24 | 3,24 | 2,49 | 4,39 | 3,23 | 2,75 | 4,11 | 3,07 | 2,92 | 5,00 | 3,46 | 2,77 |
| 31 | 7,49 | 5,29 | 4,51 | 4,18 | 3,10 | 2,47 | 3,51 | 2,26 | 2,14 | 4,57 | 3,05 | 2,28 |
| 32 | 3,64 | 2,37 | 1,96 | 2,42 | 1,36 | 1,15 | 2,16 | 1,12 | 1,02 | 2,98 | 1,76 | 1,40 |
| Average | 4,79 | 3,69 | 2,83 | 3,85 | 2,89 | 2,26 | 3,52 | 2,58 | 2,18 | 4,21 | 2,95 | 2,35 |
| St. Dev. | 2,35 | 1,96 | 1,70 | 2,64 | 2,26 | 2,01 | 2,68 | 2,13 | 1,96 | 1,96 | 1,50 | 1,39 |

Within-session reliability of AT intra-tendinous sliding (Bottom-Up Relative displacement). ICC (intra-class coefficient of correlation) and SEM (standard error of measurement) are reported for isometric contractions at 60% MVC neutral, 30% MVC neutral, 30% MVC toes-in and 30% MVC toes-out. N represents the number of participants included in the reliability analysis per condition (i.e. participants who had at least 3 trials, on the 4 recorded, with good quality).

|  | **60% MVC NEUTRAL** | **30% MVC NEUTRAL** | **30% MVC TOES-IN** | **30% MVC TOES-OUT** |
| --- | --- | --- | --- | --- |
| N (/29) | 19 | 20 | 23 | 18 |
| ICC | **0.90** (0.80 - 0.95) | **0.81** (0.78 – 0.97) | **0.84** (0.60 – 0.95) | **0.90** (0.85 – 0.97) |
| SEM (mm) | 0.09 | 0.17 | 0.13 | 0.12 |
